# Single-cell lineage tracing by endogenous mutations enriched in transposase accessible mitochondrial DNA

**DOI:** 10.1101/480202

**Authors:** Jin Xu, Kevin Nuno, Ulrike M. Litzenburger, Yanyan Qi, M Ryan Corces, Ravindra Majeti, Howard Y. Chang

## Abstract

Simultaneous measurement of cell lineage and cell fates is a longstanding goal in biomedicine. Here we describe EMBLEM, a strategy to track cell lineage using endogenous mitochondrial DNA variants in ATAC-seq data. We show that somatic mutations in mitochondrial DNA can reconstruct cell lineage relationships at single cell resolution with high sensitivity and specificity. Using EMBLEM, we define the genetic and epigenomic clonal evolution of hematopoietic stem cells and their progenies in patients with acute myeloid leukemia. EMBLEM extends lineage tracing to any eukaryotic organism without genetic engineering.

## Main Text

Resolving lineage relationships between cells is necessary to understand the fundamental mechanisms underlying normal development and the progression of disease. In recent years, new methods have emerged to enable cell lineage tracking with increasing resolution, leading to substantial biological insights^1^. Specifically, genome editing of reporter constructs via CRISPR-Cas9 allowed synthetic reconstruction of cell lineage relationships in model organisms, and has been coupled with transcriptome profiling to inform cell fates^2^. These prospective “mutate-and-record” methods provide powerful tools to resolve the developmental origin of cells in genetically engineered cells and organisms, but cannot be utilized in living humans, archival clinical samples, or any wild type organism^1^. Given these limitations, retrospective lineage tracing using endogenous genetic markers is an alternative solution. Recent advances in sequencing enable naturally occurring somatic mutations to be used as lineage markers, which usually required single-cell genome sequencing to capture the sparse genetic information^3,4^. Regions with high mutation rates, such as microsatellite repeats, retrotransposons, and copy-number variants, has been used to resolve the lineage relationship for normal or cancerous tissue samples^5,6^. These methods reduce the cost of whole genome sequencing, but still lack information on cell phenotypes. Simultaneous measurement of the lineage relationship and cell fates is ultimately required to address many biomedical questions. Here we describe EMBLEM (<u>E</u>pigenome and <u>M</u>itochondrial <u>B</u>arcode of <u>L</u>ineage from <u>E</u>ndogenous <u>M</u>utations), a strategy to track cell lineage using endogenous mitochondrial DNA variants in ATAC-seq data. The end result of EMBLEM is single-cell lineage information and rich global epigenomic profile from the same individual cells (**Fig.1a and Supplementary Figs. 1**).

**Figure 1.**
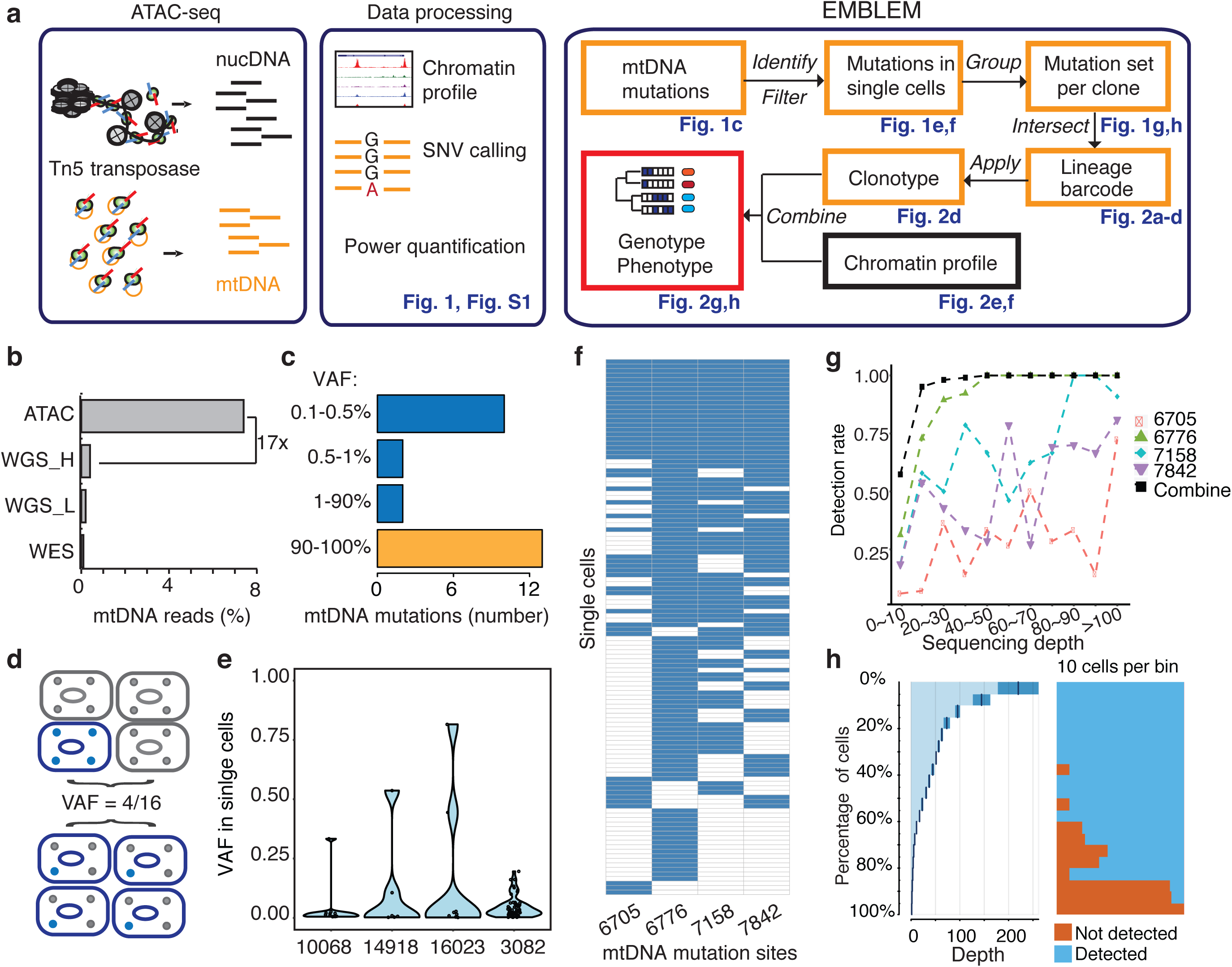
EMBLEM reveals cell lineage from mtDNA mutations. (a)EMBLEM workflow. Usings standard ATAC-seq data as input (left), an SNV calling step was added to enumerate all single nucleotide variants in mtDNA (middle). EMBLEM identifies heteroplasmic mtDNA mutations in single cells, groups mutations into diagnostic sets, and infers cell lineage based on mtDNA variants, and overlays clonotype information on epigenomic profile of the same cells (right). (b)ATAC-seq enriches for mtDNA reads compared to whole exome sequencing (WES), low coverage whole genome sequence (WGS_L), or PCR-free, high-coverage whole genome sequence (WGS_H). (c)Bimodal distribution of variant allele frequency (VAF) of mtDNA mutations discovered using ATAC-seq. Yellow bar presents the homoplastic variants that can distinguish different individuals. Heteroplasmic variants can distinguish clonal cell populations within one indiviual. (d)Two possible models for 25% mtDNA VAF in bulk: Homoplastic variants in a small proportion of cells (top) or heteroplasmic variants in nearly every single cell (bottom). Blue cells: cells with mutated mtDNA, blue dots: mtDNA with mutated allele. (e)VAF of mtDNA mutations in single cell ATAC-seq data of human B cells. Each dot present the VAF (y-axis) in single cells, and rotated kernel density on each side present their distribution. The x-axis indicates the mutation site (the nucleotide position in mitochondrial genome). (f)mtDNA mutations in human AML. Each row in the heap map is a single cell (LSC or AML blast); each column is a heteroplasmic mtDNA mutation. Blue color indicates the mtDNA variant is detected (>1 reads); white color indicates no mutation. The nucleotide position in mitochondrial genome for each mutation is indicated. (g)Combined set of heteroplasmic mtDNA mutations improve cell lineage assignment in single cells. Cells were first separated into bins according to their mtDNA coverage (x-axis). The detection rate (y-axis) for each site (indicated by different color and shape) is calculated with the number of cells with that mutations divided by total number of cells in that bin. The detection rate of combining four sites (black line, METHOD) is substantially increased. (h)Quantitation of mtDNA mutation detection rate as a function of sequencing depth and number of single cells. Cells were sorted in descending order by their sequencing depth and grouped into bins (10% of cells in each row). Distribution of sequencing depth is shown on the left panel. Cells with or without mtDNA variants are shown in blue and orange, respectively.

Assay of Transposase-Accessible Chromatin by sequencing (ATAC-seq) is a sensitive method used to study chromatin accessibility profiles in diverse cell types and organisms^7^. During DNA transposition and amplification in cells, mitochondrial DNA is also amplified at the same time (**Fig. 1a**). Mitochondrial DNA (mtDNA) is a ∼16kb circular genome with ∼10-fold higher mutation rate compared to the nuclear genome. Hence, mtDNA incrementally accumulates unique, irreversible genetic mutations that are passed on to daughter cells even in healthy humans and may be used for lineage tracing^8,9^. The majority of somatic mtDNA mutations are noncoding and thought to be passenger^10^. Importantly, the number of mitochondria (and therefore mtDNA) range from several hundreds to >10,000 per cell in different cell types, facilitating robust mtDNA analysis even from a single cell.

We first observed that ATAC-seq effectively enriches for mtDNA. While mtDNA is present in many kinds of DNA sequence libraries, it is substantially enriched in ATAC-seq libraries due to the fact that mtDNA is not chromatinized and is therefore highly accessible. ATAC-seq enables a 17-fold or greater enrichment of mtDNA compared to exome sequencing or whole genome sequencing in GM12878 human B cells (**Fig. 1b**), leading to an average ∼18,000X coverage of mtDNA (**Supplementary Figs. 2a**). With this coverage, we detected 27 mitochondrial variants from GM12878 cells (**Fig. 1c**). 13 of these variants have a variant allele frequency (VAF) greater than 90%, which are known as homoplasmic variants (**Supplementary Figs. 2b**). We also detected 14 low frequency mitochondrial DNA variants, with VAFs ranging from 0.1% to 24% (**Fig. 1c**).

The VAF from bulk ATAC-seq data represents the average of the allele frequencies of the cell population. A 25% VAF may arise from 25% of cells in the population with a homoplasmic variant, or alternatively arise from 100% of cells all having a quarter of their mitochondria with the variant allele (**Fig. 1d**). To distinguish between these two models, we analyzed single-cell ATAC-seq data from GM12878. For 4 mtDNA variants (VAF between 0.5%∼24% at population level), we find that a mixture of both models is in action for different variants (**Fig. 1e**). For instance, mtDNA mutation 3082 is widely spread among single cells, but at low frequency per cell. Because it is extremely unlikely (see METHOD) that the identical mutation arose independently in every single cell, cells sharing the same mitochondrial mutations are inferred to have descended from the same ancestral cell. These results suggest that even low frequency heteroplasmic mtDNA mutations can be exploited for lineage tracing.

To prove the principle that somatic mitochondrial mutations can track cells from the same ancestor and to quantify the power of this technique, we next applied EMBLEM to primary blood cells from patients with acute myeloid leukemia (AML). Human AML is organized as a hierarchy: A hematopoietic stem cell suffers an initiating mutation in a small number of chromatin modifier genes, and is termed the pre-leukemic hematopoietic stem cell (pHSC). pHSCs are not able transplant AML, but upon accumulation of additional mutations, they give rise to leukemic stem cells (LSCs) that are able to self-renew and recapitulate AML disease upon transplantation^11,12^. Finally, LSCs give rise to the bulk leukemic blast cells in AML^12^. Targeted exome sequencing in these same samples have identified clonal mutations in tumor suppressor genes and oncogenes that link the lineage relationship of LSCs and blasts, providing the ground truth for our analyses^13^.

We applied EMBLEM to the ATAC-seq profiles of FACS-purified pHSC, LSCs and leukemic blasts that we previously reported^14^. We first detected heteroplasmic variants from bulk ATAC-seq and then investigated whether the variants exist in each single cell using single-cell ATAC-seq. We found the LSC and blast populations not only shared the same heteroplasmic variants, but also showed similar distribution and allele frequency at the cellular level (**Supplementary Figs. 3**). These results indicate the two population are identical at the genetic level, but divergent at the epigenomic level which has been shown in previous studies^14,15^. In patient SU353, we identified four diagnostic mtDNA mutations in the same cell **(Fig. 1f)**, which indicates these four mitochondrial variants already co-existed in the ancestral cell (see METHOD). With the assumption that all these LSCs and blasts are clonal, we further quantified the detection rate of each mtDNA variant as a function of allele frequency and sequencing depth (**Fig. 1g**). We found that when a single variant allele has a frequency greater than 20%, the detection rate can be up to 90% with >20X coverage (e.g. site 6776). In contrast, when the variant allele has a frequency lower than 1%, the detection rate drops to 20% when the coverage is below 100X (e.g. site 6705). While high drop-out rate is a common challenge for single-cell technologies^16^, computational imputation of the missing information from single cell data can address this problem^17^. When multiple mtDNA variants are co-detected in multiple single cells, we can infer their origin and linkage in the ancestral cell (see METHOD). Thus, cells containing any one of these variants will still inform their origin from the same lineage. With any combination of the four variants, 90% (sensitivity) of the cells can be unambiguously assigned to the correct lineage with just 20x mtDNA coverage (**Fig. 1h**). Similar performance of single cell lineage tracing for another patient (SU070) are shown in **Supplementary Figs. 4.** These results demonstrate that somatic DNA mutations in the mitochondrial genome are a powerful endogenous marker to identify clonal cell populations.

To expand on these findings to additional different cell lineages, we applied EMBLEM to bulk ATAC-seq data from sorted blood cells from 9 healthy human donors and 11 patients with AML^14^. We identified homoplasmic and heteroplasmic mtDNA mutations in multiple lineages of primary blood cells from healthy donors and all AML patients (**Fig. 2a and Supplementary Figs. 5a**). The homoplasmic variants from each patient depicted the phylogenetic relationship of individuals, permitting identification of cells from individual patients on the basis of mtDNA variants (**Fig. 2a**). The heteroplasmic mutations shows a similar mutant spectrum as observed by previous studies using cancer genomic data **(Supplementary Figs. 5b and c**)^10^. Furthermore, the heteroplasmic mtDNA mutations within each patient revealed the lineage relationship between FACS-sorted cell populations (**Fig. 2b-c**). In all AML cases with LSCs examined(6 cases), the LSCs and their corresponding leukemic blasts have nearly identical heteroplasmic mtDNA mutations (**Supplementary Figs. 5d-i**), suggesting a direct lineage relationship and short generation history between LSCs and blasts. We then examined whether any of the mtDNA variants present in LSCs can be seen in the pHSCs, where the first leukemia-associated protein-coding mutations have already occurred in functional normal hematopoietic stem cells^13,14^. We detected LSC-associated mtDNA mutations in pHSCs in all 11 cases. Interestingly in 7 of the 11 cases, we also detected additional heteroplasmic mtDNA mutations present only in pHSCs (**Supplementary Figs. 5g-h**). pHSCs are capable of long-term self-renewal and possess a clonal growth advantage, allowing them to clonally outcompete normal HSCs. Indeed, the clonal frequency of pHSCs is a poor prognostic factor for overall survival in AML^14^. Our discovery of pHSCs with distinct heteroplasmic mtDNA mutations suggests the existence of multiple distinct sub-clones of pHSCs in AML patients.

**Figure 2.**
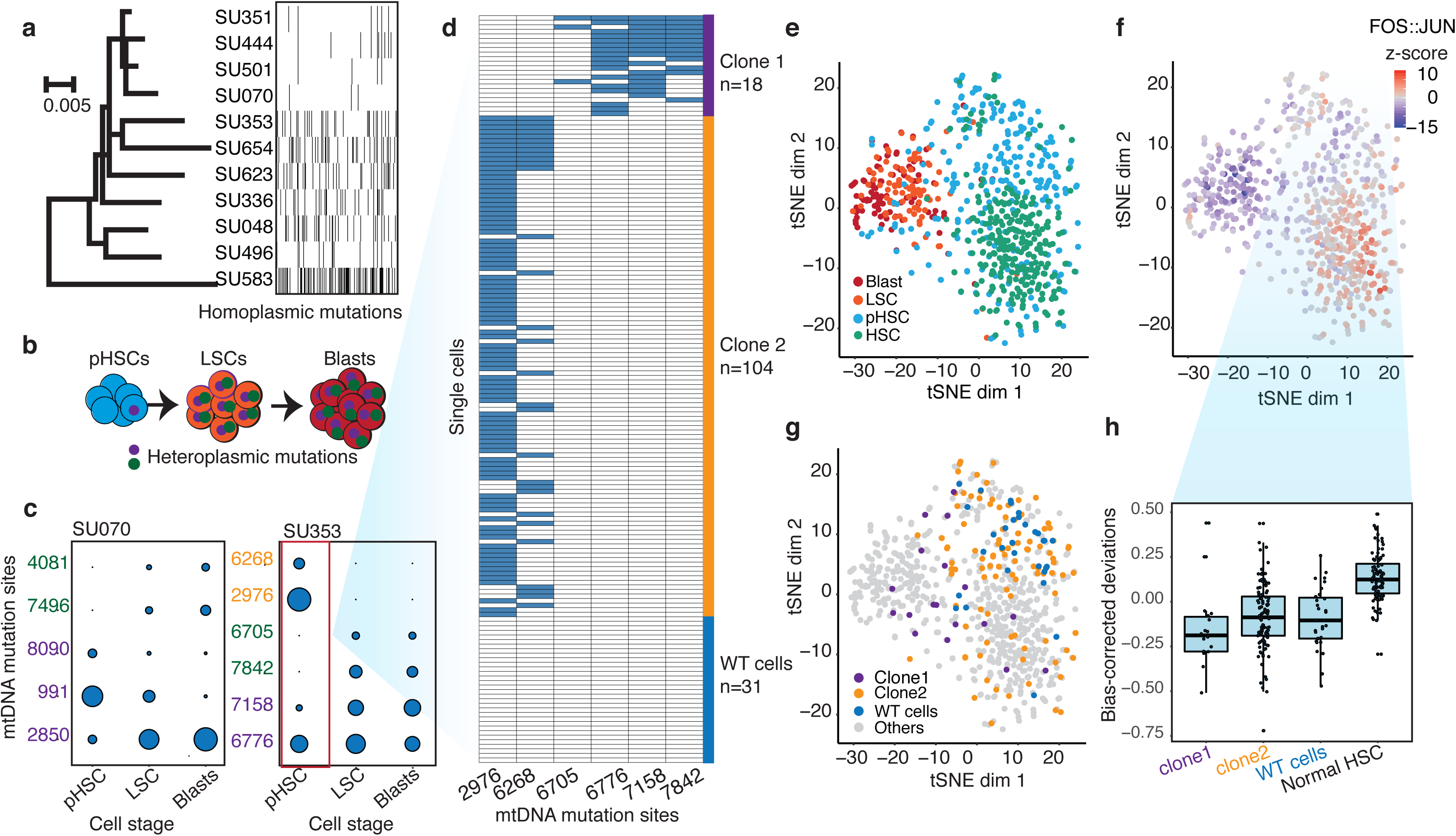
Clonal evolution of pre-leukemic HSCs inferred from joint lineage tracing and single cell chromatin accessibility. (a)Personal identification of blood cells from healthy donors and patients using homoplastic mtDNA variants. The black dashes represent homoplasmic mtDNA variants in each individual. The tree depicts the polygenetic relationship among the individuals; the branch lengths depict the number of substitutions per site with scale bar shown. (b)Scheme of using heteroplasmic mtDNA mutations to track cell lineages during AML development. (c)EMBLEM deconvolutes AML clonal heterogeneity. Heteroplasmic mtDNA mutations in three cell populations from two AML patients are shown (left panel). Mutations sites (in rows) in each FACS-sorted cell population (in columns) are shown, with size of each circle representing its VAF. Several mtDNA mutations (sites shown in purple) are detected in pHSCs and transmitted to LSCs and blasts, confirming those pHSC clones at the apex of leukemia lineage. LSCs accumulated additional mtDNA mutations (sites shown in green) and are transmitted to leukemic blasts in patient SU070. In patient SU353, in addition to shared mtDNA mutations in pHSCs, LSCs, and blasts (purple), two pHSCs-specific mtDNA mutations are also detected (yellow). Detail information of VAF and sequencing depth of variant allele are shown in **Supplementary Figs. 3 and 4** (d)Heteroplasmic mutations in single pHSCs from one patient reveals clonal heterogeneity. Each column is a mtDNA nucleotide position; each row is one cell. Blue color indicates the presence of the mtDNA variant. Shown are cells with any mtDNA mutation detected, or cells with more than 40X coverage of the mitochondrial genome without any detected mutatino. The number of cells in each clonotype are indicated on the right. (e)Landscape of single-cell chromatin accessibility of blood progenitor and leukemic cells in patient SU353. tSNE map using bias-corrected deviations from chromatin accessibility showing cluster of AML blasts, LSCs, pHSCs and normal HSC, colored by cell types. (f)Chromatin accessibility of the FOS:JUN binding motif across the same single cells. tSNE map colored by deviation z-score for motif associated to FOS:JUN, the most variable TF motif. (g)pHSC clones possess distinct epigenomic signatures. Clone 1 that gives rise to the AML has a chromatin accessibility profile that more resembles LSCs and leukemic blasts. Clonotype information from EMBLEM is overaid on the tSNE map defined by TF motif deviations, and colored by different lineal sub-populations defined by mtDNA mutations. (h)Quantitation of distinct single-cell chromatin accessibility at FOS:JUN motifs among different pHSC clones defined by EMBLE. Clone 1 pHSCs tend to down regulate FOS:JUN accessibility, while clone 2 pHSC shows substantially greater cell-to-cell variability. pHSCs with no detectable mtDNA variants and normal HSCs are shown for comparison. TF deviation of single cells (black dots) is shown on the distribution box-plot.

To validate the heterogeneity of pHSCs inferred from EMBLEM of bulk cell populations, we performed single-cell ATAC-seq of HSCs from AML patient SU353, which exhibited both a high burden of pre-leukemic somatic coding gene mutations and high frequency of pHSC-specific heteroplasmic mtDNA mutations^14^. We identified the heteroplasmic mtDNA variants from each single cell, which separated the HSCs into three lineages: Two clonal subpopulations termed “clone 1” (18 cells) and “clone 2” (104 cells), and a third population with no mtDNA variants despite sufficient mtDNA coverage (WT, 31 cells) (**Fig. 2d**). Notably, clone 2 possessed pHSC-specific mtDNA mutations, while clone 1 possessed mtDNA mutations shared with LSCs, indicating clone1 is the lineage precursor of AML. These results confirm that multiple pHSC clones arise in AML patients, and one subclone eventually evolved to become the LSC.

Finally, we related the clonotype of pHSCs to their single-cell chromatin accessibility profiles. We interrogated the patterns of active DNA elements and enriched transcription factor motifs in sequential stages of AML development from the same patient, and contrasted with HSCs from normal donors using ChromVAR^15^ (**Fig. 2e and Supplementary Figs. 6a**). The chromatin accessibility profiles of pHSCs are more similar to HSCs than to LSCs or leukemic blasts. The greatest deviation between HSC and other cell types occurred at DNA binding motifs of the transcription factor Jun/Fos, a known key regulator of HSC biology^18^ (**Fig. 2f**). Furthermore, the three lineages of pHSCs revealed by mtDNA mutations also showed distinctive chromatin profiles (**Fig. 2g**). Clone 1 pHSC, which gives rise to the LSC and AML leukemia, is already more similar to LSCs and blasts in its chromatin accessibility. In contrast, clone 2 that comprises the larger fraction of pHSCs exhibited variable chromatin profiles at the single-cell level that spanned the range of normal HSCs. Thus both lineage tracing and single cell epigenomic states suggest clone 1 as the original stem cell of the AML in patient SU353. Supervised comparison of the chromatin accessibility profiles among these clonal sub-populations further identified distinct and significantly enriched transcription factor motifs (**Fig. 2h, Supplementary Figs.6b-d**). These results indicate the heterogeneity of HSCs from AML patients both on a genetic and epigenomic level. This pilot study provides a practical approach to clarify the process of pre-leukemic clonal evolution and informs the emerging biology of clonal hematopoiesis.

In summary, we present a computational strategy by which we can trace cell lineage by endogenous mtDNA mutations and combine chromatin accessibility profiling in the same cell using ATAC-seq data. This approach is applicable to any eukaryote, does not require genetic engineering or genome editing, and is cost effective as the lineage information comes “for free” on top of epigenomic insights. EMBLEM may also be extended to other single cell technologies, in which mtDNA is sequenced. We show that EMBLEM is successful even with low frequency heteroplasmic mutations, detection of rare clones in a population, and authentic clinical samples. Potential limitations to this approach include instances of biased mtDNA replication and segregation and rare inter-cellular transfer of mitochondria^19^. With advances in the throughput and depth of single-cell genomic technologies, we believe EMBLEM may be a powerful tool to bring insight for many biomedical questions, including development, regeneration, immunity, and cancer.

## Methods

Methods, including statement of data availability and any associated accession code and reference, as available online.

## Supporting information

## Author contributions

J.X. and H.Y.C conceived the project. J.X. performed the data analysis, and J.X, K.N., U.M.L, Y.Q. performed the experiments and M.R.C. provided the resequencing data. R.M and K.N provided the samples for this study. H.Y.C. and R.M. supervised the project and wrote the paper with J.X and K.N.

## Acknowledgements

We thank C. Curtis, Ava Carter, Furqan Fazal, Kevin Parker and Chun-Kan Chen for insightful advice and assistance. Supported by US National Institutes of Health P50-HG007735 (to H.Y.C.), R01CA188055 (to R.M.), and R01HL142637 (to R.M.). H.Y.C. is an Investigator of the Howard Hughes Medical Institute.

## Competing interests

H.Y.C. is a co-founder of Accent Therapeutics and an advisor for 10x Genomics and Spring Discovery. Stanford University has filed a patent on ATAC-seq, on which H.Y.C. is named as an inventor.

