## Supplementary figures and images for "Single-cell lineage tracing by endogenous mutations enriched in transposase accessible mitochondrial DNA"

### Supplementary file 1

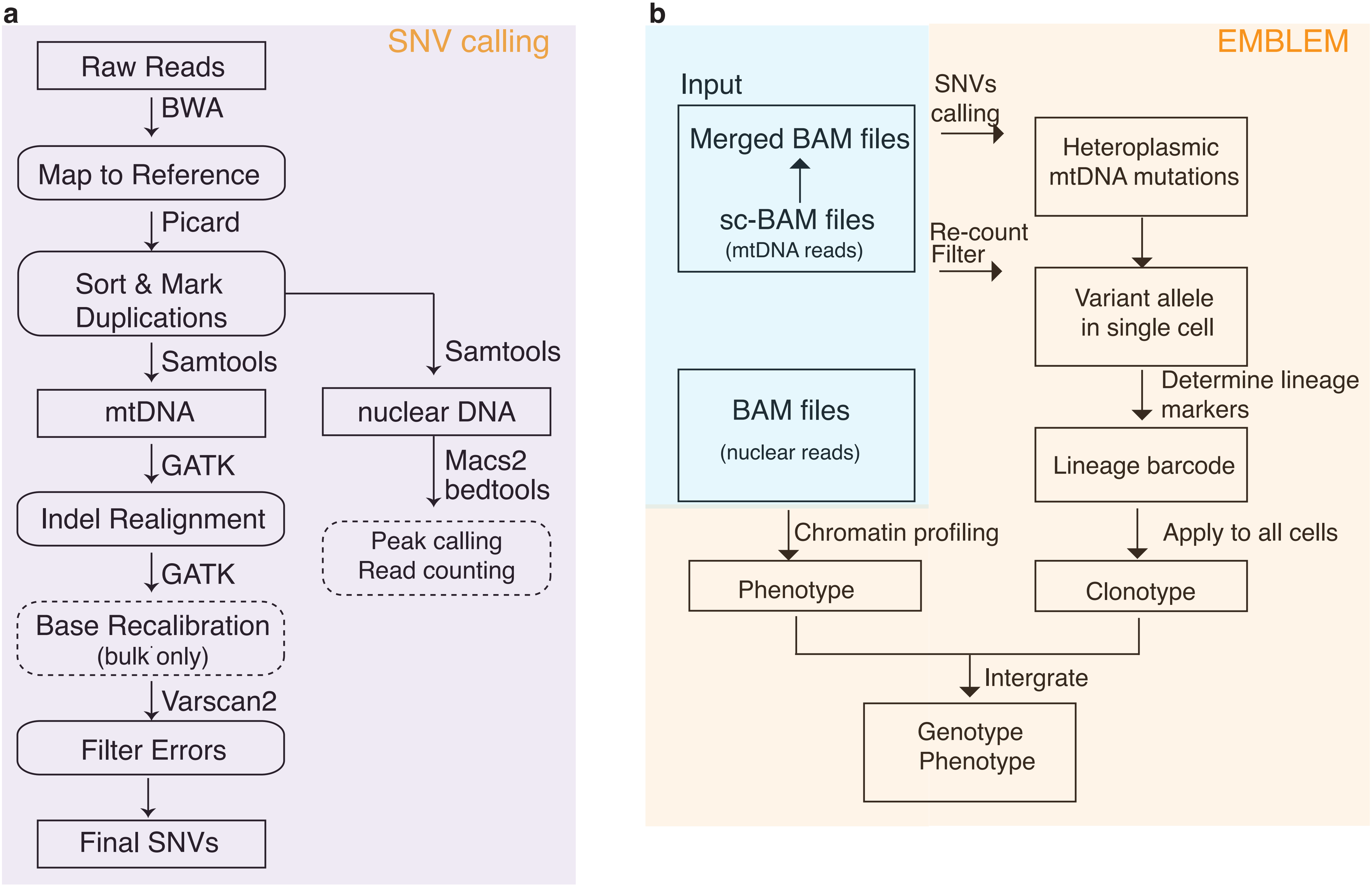

### Supplementary file 2

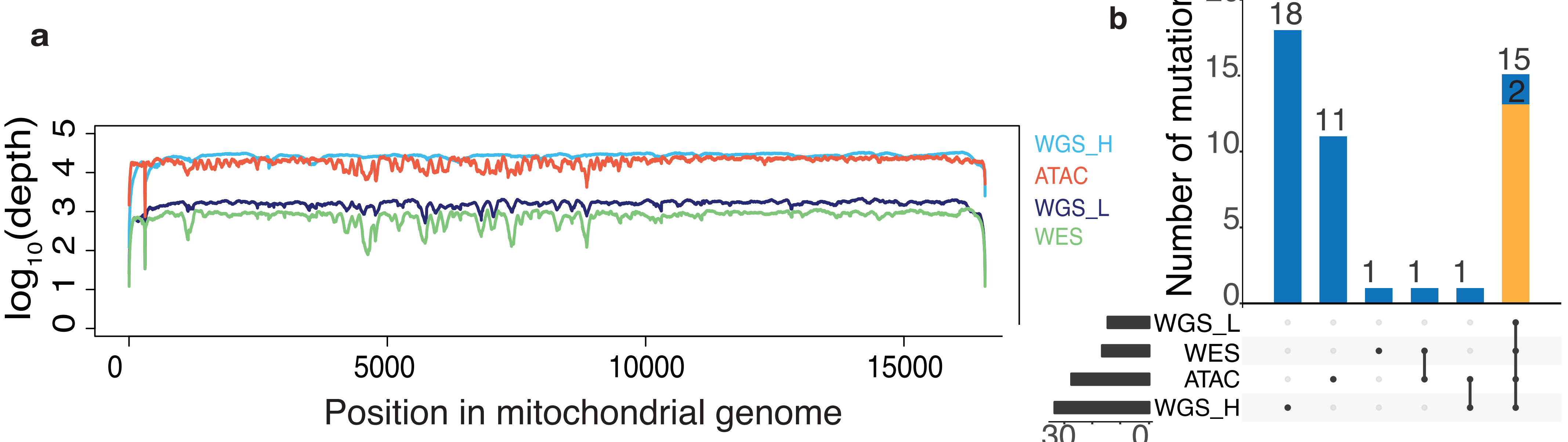

### Supplementary file 3

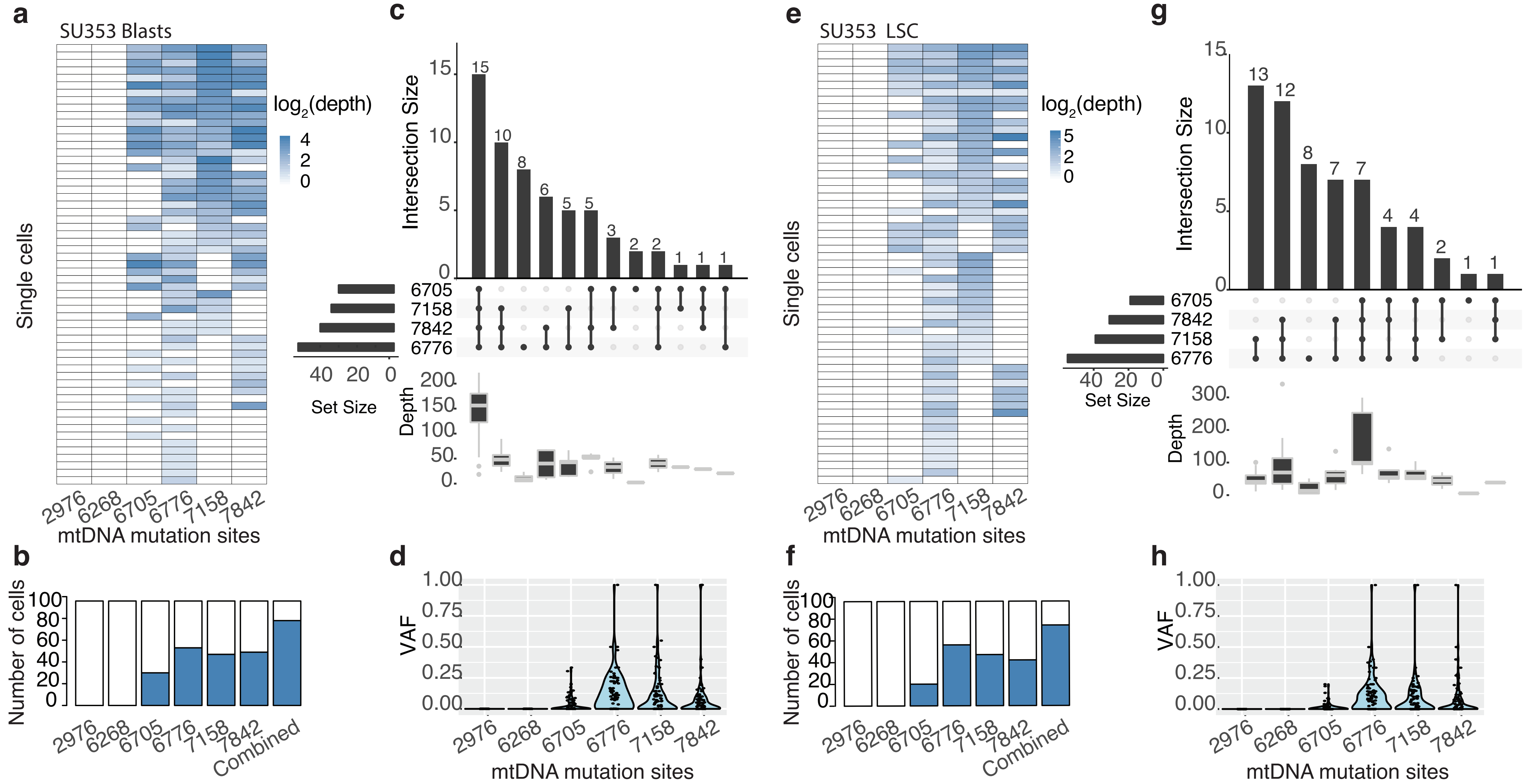

### Supplementary file 4

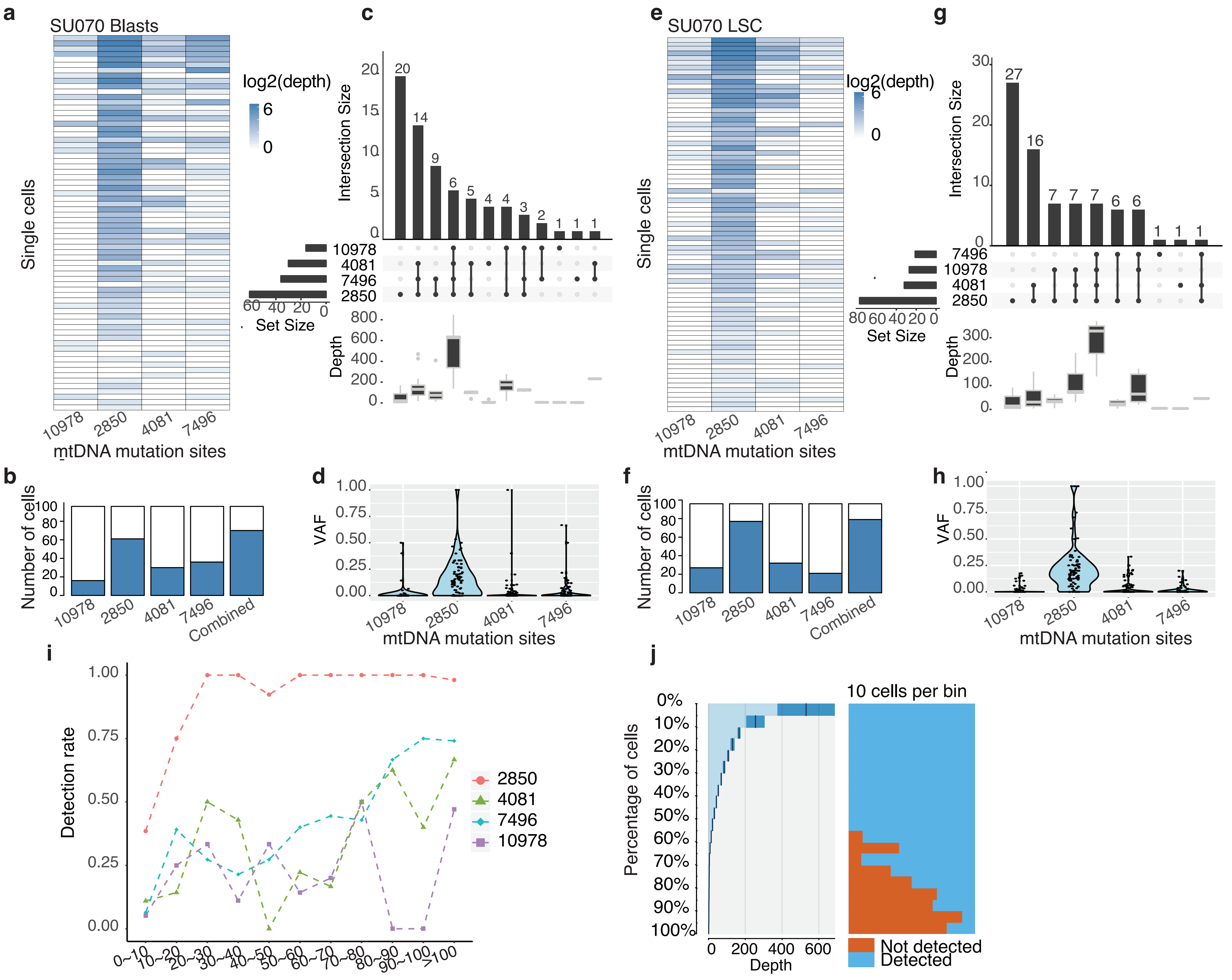

### Supplementary file 5

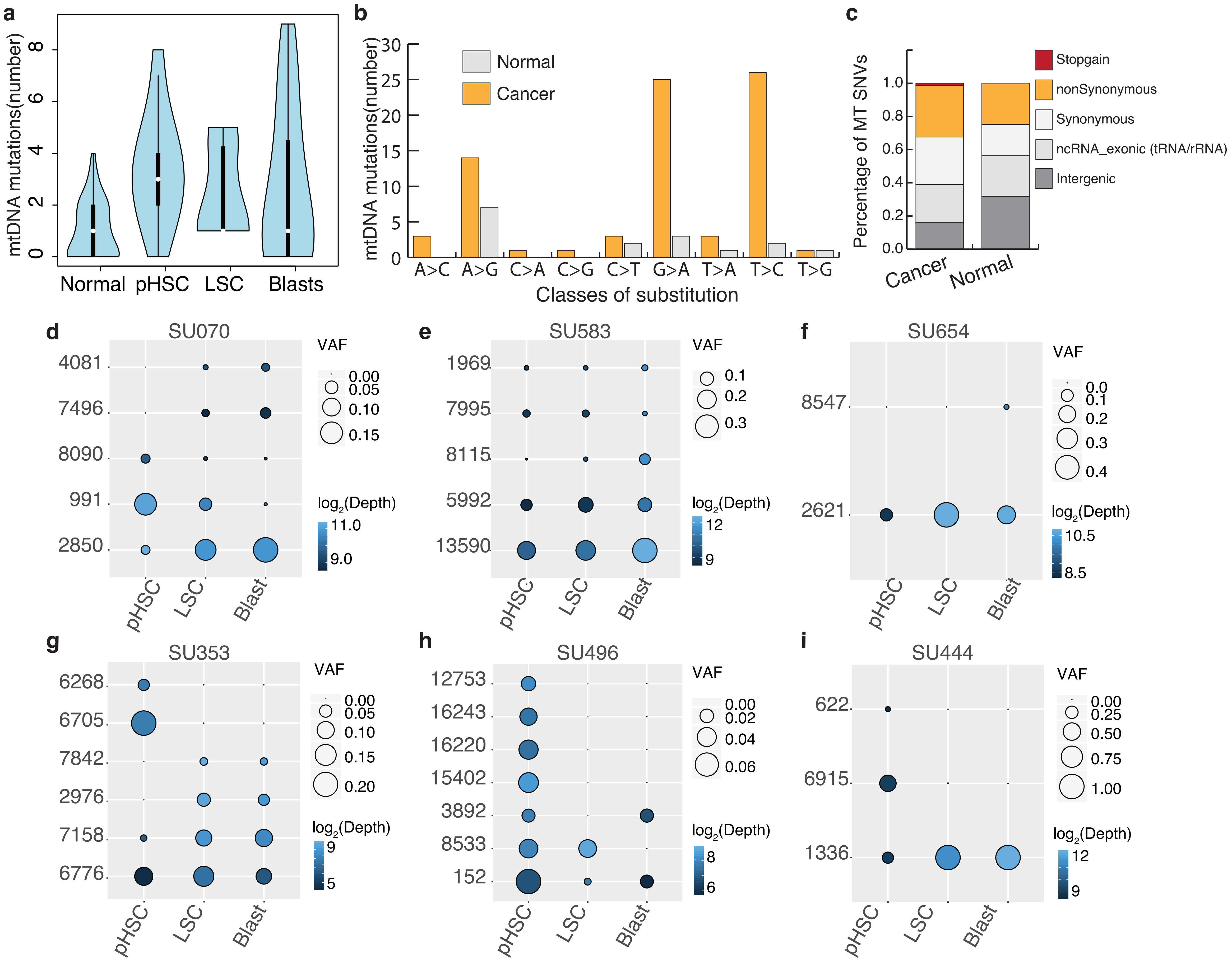

### Supplementary file 6

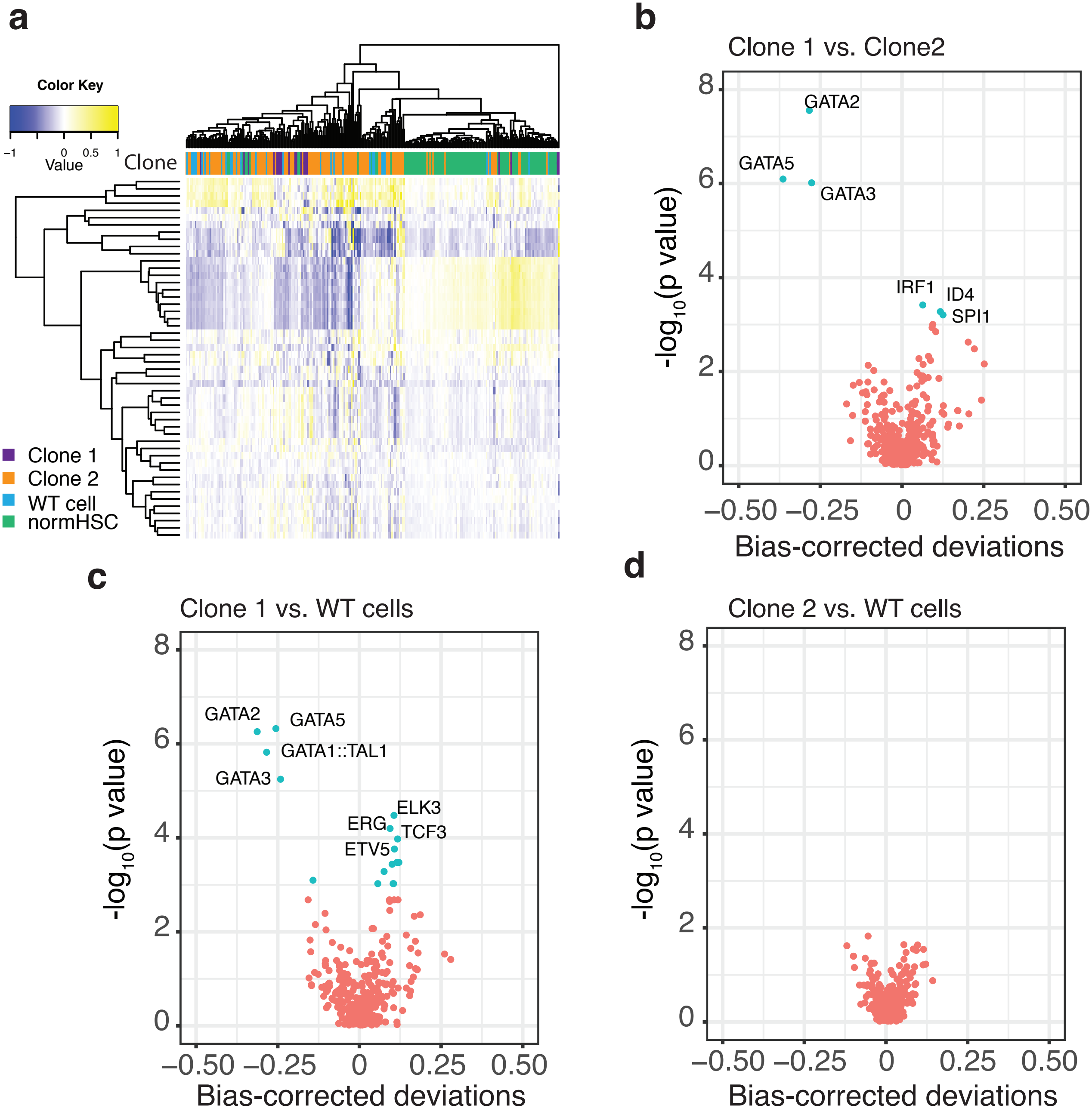

### Supplementary file 7

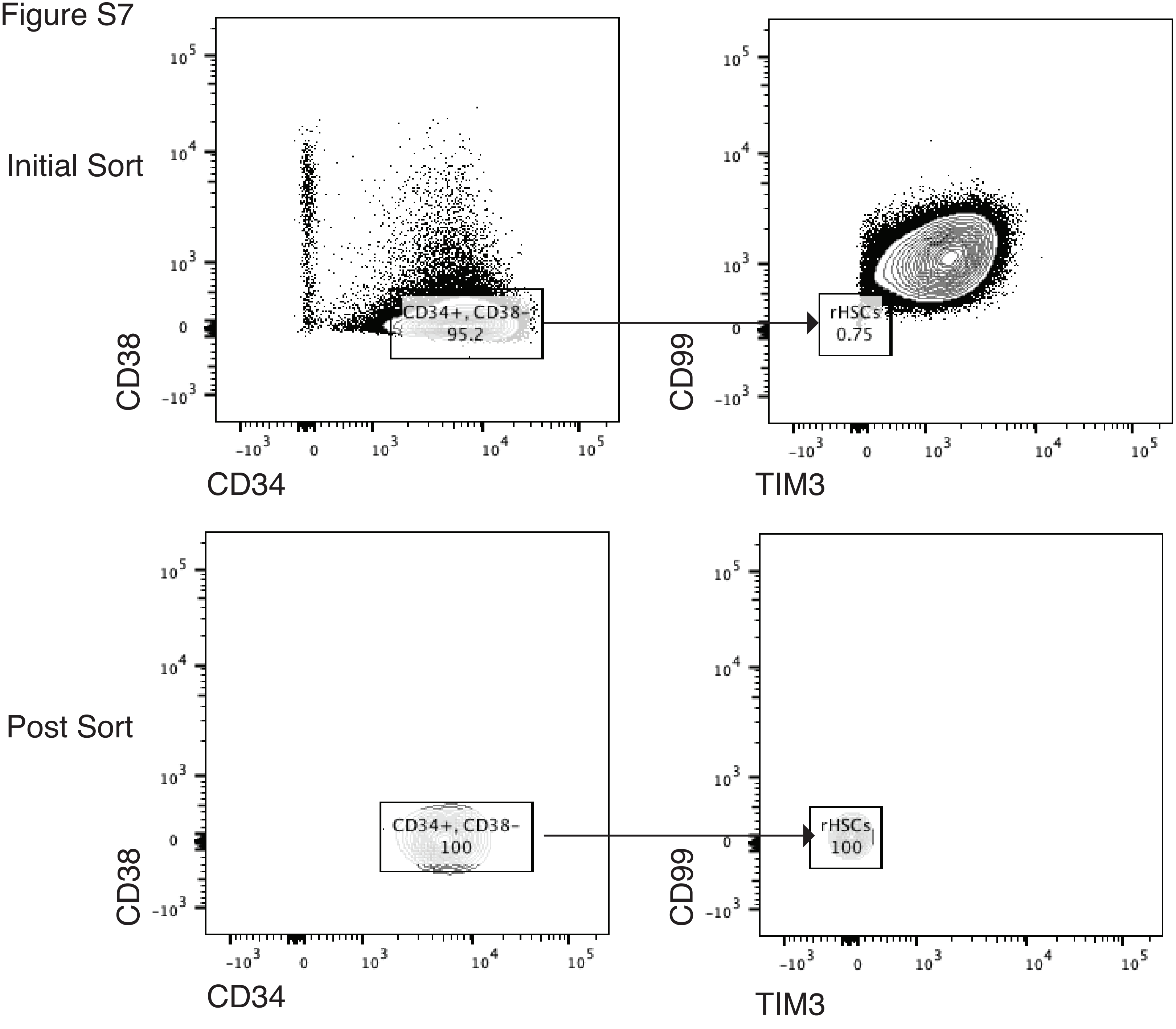

### Supplementary file 8

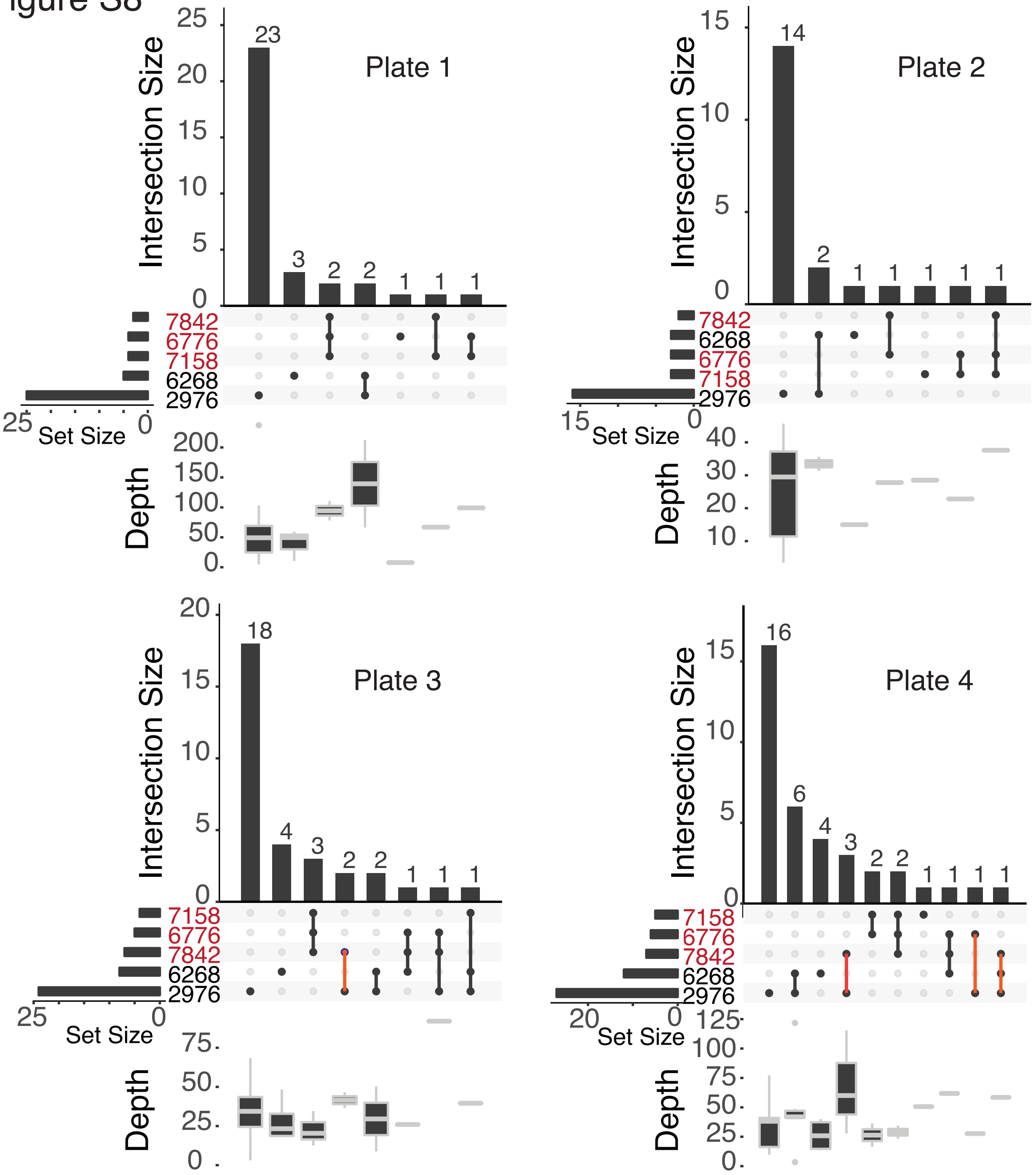
