## Supplementary material for "Single-cell lineage tracing by endogenous mutations enriched in transposase accessible mitochondrial DNA"

**Supplementary figure legends**

**Figure S1** **EMBLEM workflow for SNP calling and lineage inference.**

(**a**) Workflow for mitochondrial DNA variant calling from ATAC-seq data. This workflow was applied to both bulk and single cell ATAC-seq. The steps indicated with dotted lines were not applied to single-cell data.

(**b**) Workflow for inferring lineage relationships from single-cell ATAC-seq data. BAM files from single cells were first merged and confident mtDNA variants were called. Mutated alleles from these variant sites were then counted for each single cell. The cell lineage was then inferred from mtDNA variants and analyzed alongside the chromatin profile for each cell.

**Figure S2 mtDNA coverage and variants from different sequencing libraries from GM12878 human B cells.**

(**a**) Mitochondrial genome coverage from each of four different sequencing libraries including WGS_H (high coverage PCR-free whole genome sequencing), WGS_L(low coverage whole genome sequencing), WES(whole exome sequencing), ATAC-seq. The Y axis shows coverage scaled in log_10_. 43M paired-end ATAC-seq reads(2x50bp) yielded the same coverage of mtDNA as 747M paired-end reads(2x250bp) from WGS-H data.

(**b**) Comparison of variants detected in sequencing data from four different library preparations. The number of variants detected in each library is shown on the bottom left. The intersection of different libraries (bottom-right) and the number of variants s are shown on the top. Homoplasmic variants are in yellow and heteroplasmic variants are in blue.

**Figure S3** **Heteroplasmic variants in single cells from AML blasts and LSCs (SU353)**

(**a**) Heatmap showing variant mitochrondrial sites (columns) in each AML blast from patient SU353(rows). The color represents the number of reads supporting the variant allele (log2(depth)). The first two sites are negative controls, which are detected in pHSCs only.

(**b**) Bar plot showing the number of cells in which we detect each mitochondrial variant. The last bar shows the number of cells with any one of the four variants detected.

(**c**) The top right shows the number of cells with each different combination of variants detected. The number of cells is shown on top of the bar. The combination of variants detected is annotated below the bar. The total number of cells with each variant site detected is shown to the left. The average coverage of the mitochondrial genome for each intersection group is shown below.

(**d**) VAF of mtDNA variants.The x-axis indicates the variant site notated by the nucleotide position in the mitochondrial genome. Each dot represents the VAF (y-axis) in single cells and the rotated kernel density on each side shows their distribution.

(**e-h**) Same as (**a-d**), for leukemia stem cells (LSCs) from patient SU353.

**Figure S4 Heteroplasmic variants in single cells from AML blasts and LSCs (SU070)**

(**a-d**) Same as Figure S3 a-d, for AML blasts from SU070.

(**e-h**) Same as Figure S3 e-h, for LSCs from SU070.

(**i**) Quantification of the detection rate for each heteroplasmic variant from mtDNA. Cells (both LSCs and AML blasts) were first separated into bins according to their coverage of mtDNA (x-axis). The detection rate (y-axis) for each site (notated by different color and shape) is calculated as the number of cells with that variant detected divided by the total number of cells in that bin.

(**j**) Quantitation of mtDNA mutation detection rate as a function of sequencing depth and the number of single cells. Cells were sorted in descending order by their sequencing depth and grouped into bins (10 cells in each row). Distribution of sequencing depth is shown on the left panel. Cells with or without mtDNA mutations are shown in blue or orange, respectively.

**Figure S5 Heteroplasmic variants in AML patients.**

(**a**) The number of heteroplasmic variants detected using ATAC-seq data from normal primary blood cells and cancer cells from AML patients.

(**b**) The number of mtDNA variants identified from normal and cancer samples in different substitution classes are shown as a bar plot. Mutations from normal (gray) and cancer (yellow) samples are separated. The C>T and T>C signature in cancer mtDNA has been observed in previous studies and it's equivalent to the one that has been operating during the evolution of human germline mtDNAs.

(**c**) Annotation of mtDNA mutations and the proportion of mutations in coding and non-coding regions. Coding mutations are divided into synonymous, nonsynonymous, and gain of stop codon. Heteroplasmic mutations detected from cancer samples show a similar distribution as those from normal samples, with a slightly higher proportion falling within coding regions.

(**d-i**) Heteroplasmic mutations in three cell stages for each AML patient. Variant allele (in rows) in each cell population (in columns) are shown with a circle, with size indicating their variant allele frequency. Sequencing depth of the variant allele is indicated by the color of the circle (in log_2_ scale).

**Figure S6 Single cell chromatin accessibility**

(**a**) Heat map showing clustering of pHSCs from SU353 and normal HSCs from a healthy donor, based on the z-score of TF deviation. The Z-scored deviation is shown for individual cells (columns) for each TF (rows). Clone information is shown on the top of the heat map. Top 50 most variable motifs were used in this heat map.

(**b**) Volcano plot showing the difference in chromatin accessibility for transcription factor binding motifs between Clone 1 and Clone 2. The x-axis shows the mean difference of bias-corrected deviations and the y-axis shows the p-value (in log_10_ scale). The most significant differential motifs are annotated with TF names.

(**c**) Same as in (b) for Clone 1 vs. WT cells.

(**d**)Same as in (b) for Clone 2 vs. WT cells. No significantly differential motifs were detected.

**Figure S7 Sorting Scheme for pHSCs.**

Scheme of FACS sorting of the pHSC population from AML patient SU353. Initial sort (top panel) and post-sort purity (bottom panel) are shown.

**Figure S8 Intersection of mtDNA variants for individual C1 chip.**

The intersection of mtDNA variants in single pHSCs from four C1 chips. For each plate, the number of cells with each combination detected is shown as a bar (top panel) and the annotation of sites is shown below the bar. The total number of cells with each site detected is shown to the left. The average sequencing depth of the mitochondrial genome for each intersection group is shown under the annotations. The variants in red indicate common mutations and black indicates pHSC specific mutations.
