## Supplementary material for "Single-cell lineage tracing by endogenous mutations enriched in transposase accessible mitochondrial DNA"

**ONLINE METHODS**

**Public data accession.** Aligned bam files for GM12878 whole exome, low coverage whole genome, and PCR free whole genome sequence, were downloaded through phase 3 release of 1000 genomes (ftp://ftp.1000genomes.ebi.ac.uk)

The alignment files were accessed via the following ftp links:

- ftp://ftp.1000genomes.ebi.ac.uk/vol1/ftp/phase3/data/NA12878/exome_alignment/NA12878.mapped.ILLUMINA.bwa.CEU.exome.20121211.bam ./
- ftp://ftp.1000genomes.ebi.ac.uk/vol1/ftp/phase3/data/NA12878/alignment/NA12878.mapped.ILLUMINA.bwa.CEU.low_coverage.20121211.bam ./
- ftp://ftp.1000genomes.ebi.ac.uk/vol1/ftp/phase3/data/NA12878/high_coverage_alignment/NA12878.mapped.ILLUMINA.bwa.CEU.high_coverage_pcr_free.20130906.bam ./

ATAC-seq and single cell ATAC-seq data for GM12878 generated by Buenrostro et al. were downloaded through GEO with accession number GSE47753 and GSE65360, respectively^1,2^. Bulk ATAC-seq data from normal donors and AML patients generated by Corces et.al^3^., were downloaded through GEO with accession number GSE74912. Single cell ATAC-seq data for leukemia stem cell and leukemic blasts generated by the same study were downloaded through GEO with accession number GSE74310. Single cell ATAC-seq from normal HSC generated by Buenrostro et al., were downloaded through GEO with accession number GSE96772^4^.

**Comparison of mitochondrial genome capture rate and coverage.** Sequencing reads from ATAC-seq were aligned to the reference genome by BWA alignment tool^5^. The same reference, GRCh37(used by 1000 genome) and human reference mtDNA sequence rCRS (revised Cambridge reference sequence), were used for ATAC-seq data processing. Samtools^6^ was used for manipulating sequence reads and calculating sequence depth. For all the data sets, the aligned reads were further filtered with mapping quality (Q >30) and PCR redundancy was removed. The percentage of reads from mitochondrial genome compared to that of the nuclear genome were calculated after all the clean-up steps. The mitochondrial genome coverage was calculated using bases with sufficient sequence quality score (q > 30). A strong depletion region around 3107 due to the sequencing error(3170N) in the reference genome was excluded in the coverage plot^7^.

**Bulk ATAC-seq data process and mitochondrial DNA variants calling.** Most of the ATAC-seq pipelines remove mtDNA during their process. To rescue the genetic information from mtDNA, we modified our ATAC-seq pipeline and added SNP calling steps, which focuses on the mitochondrial genome. Briefly, adaptor sequences were trimmed from FASTQs using custom Python scripts. Paired-end reads were aligned to the reference genome using BWA. To improve the accuracy of heteroplasmic mutation calling, we followed the somatic mutation calling guidelines from GATK^8^, with additional clean-up steps before variant calling. Reads mapped to mtDNA were extracted using Samtools^6^ from the final bam files and variants were called using VarScan2^9^ with "--min-var-freq 0.001" (**Supplementary** **Figs.1a**). The heteroplasmic variants were further filtered through the following steps to exclude potential sequencing or mapping errors:

1. Thirteen frequent false-positive variants by misalignment due to extensive level of homopolymers in rCRS and due to sequencing error in the reference genome(reported in the previous study^7^), were also observed and removed in this study. The following sites were explicitly removed:

Misalignment due to ACCCCCCCTCCCCC (rCRS 302-315)

A302C, C309T, C311T, C312T, C313T, G316C

Misalignment due to GCACACACACACC (rCRS 513-525)

C514A, A515G, A523C, C524G

Misalignment due to 3107N in rCRS (ACNTT, rCRS 3105-3109)

C3106A, T3109C, C3110A

2. Strand imbalance is a potential feature of sequencing error with various causes. To remove the potential sequence error from Illumina NextSeq (with a known high error rate at A bases) and sequence error from DAN damage(G->T, C->A)^10^, we required a VAF >1%, greater than 2 reads detected from both the forward and reverse orientation, and strand is balanced (30%<forward/(forward + reverse)<70%).

3. Variant sites with VAF>0.9, but less than 1, were counted as homoplasmic variants.

Although the germline polymorphic can be a back heteroplasmic mutations, the observation of these events is higher than expected, which implies the false positive calling due to mapping bias for non-reference allele and sequencing errors.

In the 12 AML cases from Corces et.al^3^, we found that in one patient (SU209), the number of heteroplasmic mutations (37) and their VAF are significantly higher than other patients. Most of these heteroplasmic mutations also overlapped with common variants present in the general human population (http://ftp.1000genomes.ebi.ac.uk/vol1/ftp/release/20130502/ALL.chrMT.phase3_callmom-v0_4.20130502.genotypes.vcf.gz), which indicates potential sample contamination. Therefore, this case was excluded from this study.

**Single cell ATAC library resequencing.** To better evaluate the detection rate in single cell ATAC-seq data, we re-sequenced the previous libraries(LSCs and AML blasts from SU070 and SU373) from Corces et.al^3^. The re-sequenced data were uploaded to GEO and accession number is GSE122576.

**Human AML samples** Human AML samples were obtained from patients at the Stanford Medical Center with informed consent, according to institutional review board (IRB)-approved protocols (Stanford IRB, 18329 and 6453). Mononuclear cells from each sample were isolated by Ficoll separation, resuspended in 90% FBS + 10% DMSO, and cryopreserved in liquid nitrogen. All analyses conducted here on AML cells used freshly thawed cells.

**Cell Sorting.** Cell samples were first thawed and incubated at 37°C with 200 U/mL DNase in IMDM + 10% FBS. To enrich for CD34+ cells, magnetic bead separation was performed using MACS beads (Miltenyi Biotech) according to the manufacturer’s protocol.

For cell staining and sorting, the following antibody cocktail was used with the schema shown in **Supplementary** **Figs.7**

CD34-APC, clone 581, Biolegend, at 1:50 dilution.

CD38-PE-Cy7, clone HB7, Biolegend, 1:25 dilution.

CD19-PE-Cy5, clone H1B9, BD Biosciences, 1:50 dilution

CD20-PE-Cy5, clone 2H7, BD Biosciences, 1:50 dilution

CD3-APC-Cy7, clone SK7, BD Biosciences, 1:25 dilution

CD99-FITC, clone TU12, BD Biosciences, 1:20 dilution

TIM3-PE, clone 344823, R&D Systems, 1:20 dilution

CD45-KromeOrange, clone J.33, Beckman Coulter at 1:25 dilution

Samples were sorted using a Becton Dickinson FACS Aria II. pHSCs were re-suspended and kept in cold FACS buffer containing 1 ug/mL propidium iodide prior to and after sorting. Cells were then immediately prepared for single cell ATAC-seq.

**Single cell ATAC-seq from pHSC.** Cells were washed 2 times in C1 DNA Seq Cell Wash Buffer (Fluidigm). ~10K cells were then re-suspended in 6 mL of C1 DNA Seq Cell Wash Buffer, and were combined with 4 mL of C1 Cell Suspension Reagent, 7 mL of this cell mix was loaded onto the Fluidigm IFC. Cells at a concentration of 260-380 cells/µL were then assayed using scATAC-seq as previously described^2^. Briefly, single cells were captured using the C1 Single-Cell Auto Prep IFC microfluidic chips. Cells were permeabilized and accessible fragments were captured using 20 µL of Tn5 transposition mix (1.5x TD buffer, 1.5 µL transposease (Nextera DNA Sample Prep Kit, Illumina), 1x C1 Loading Reagent with low salt (Fluidigm), and 0.15% NP40) at 30 minutes at 37°C. In a 96-well plate, 7 µL of harvested libraries were amplified in 50 µL PCR for an additional 17 cycles (1.25 µM custom Nextera dual-index PCR primers in 1x NEBnext High-Fidelity PCR Master Mix using the following PCR conditions: 72°C for 5min; 98°C for 30 s;) using the following PCR conditions: 72°C for 5min; 98°C for 30 s; and thermocycling at 98°C for 10 s, 72°C for 30 s, and 72°C for 1 min. The PCR products were pooled creating a final volume of ~4.8 mL. The pooled library was purified on a single MinElute PCR purification column (Qiagen). Libraries were quantified using qPCR prior to sequencing. The scATAC-seq libraries were sequenced by Illumina MiSeq. The sequence data was uploaded to GEO under the accession number GSE122577.

**Single cell ATAC-seq data processing and mitochondrial DNA variant calling.** Single cell ATAC-seq were processed similarly to the bulk ATAC-seq, taking each individual cell as one sample. Recalibration steps were not applied for single cell data, as the sequence depth is not sufficient to empirically adjust the quality scores. After cleaning the alignment, files from every single cell were merged and heteroplasmic variants were first called with the merged bam and filtered using the same criteria as bulk data. Heteroplasmic variants called from merged data or from bulk data were re-counted in each individual cell using Samtools with "-q 20 -Q 20". And the non-reference allele had to match the variants detected in merged or bulk data.

**Detection rate estimation.** In every single cell, if the variant allele detected in merged or bulk data were supported by any reads, it was considered positive; otherwise, it was counted as zero. A binary matrix was used to present the lineage relationship among single cells and plotted as a heat map. The intersections of the variants were quantified by the Upset R package^11^. The number of detected variants showed a correlation with sequencing depth and the number of cells with all variants (**Supplementary Figs.3 and Figs.4**) confirmed the variants already co-existed in the ancestral cell. Following this assumption, the detection rate can be measured as the proportion of cells with variants in the total number of cells. For each variant, cells were separated into different bins, increased by 10, according to the total sequencing depth at each variant. The detection rate for each variant site was then calculated in each bin. The combined detection rate was estimated by 1-(1-*R_1_*)*(1-*R_2_*)*(1-*R_3_*)*(1-*R_4_*), where *R_n_* is the detection rate for each variant.

**Lineage inference.** The probability of observing a mutation at a given site is *P_n_=n*r*, where *r* is the average mutation rate in the mitochondrial genome and *n* is the copy of mtDNAs in a single cell. *r* is estimated to be ~10^^-7^ per base^12^, n is around 100~10000 per cell^13^, so *P_n_* will be 10^^-5^~10^^-3^. The probability of N cells sharing the same mtDNA mutations, but raising independently, will be (*P_n_*)^^N^. Thus, when there are more than 3 cells in the population sharing a common mtDNA mutation, the probability of these independently occurring will be close to 0. Cells with common mtDNA mutations inherited the mutations from the same ancestral cell is more likely to explain the observation. Furthermore, when a set of mutations (more than 1) is detected in more than 1 cells, the null hypothesis (independently occurred) is rejected more confidently. The mutations within the ancestry cells can be inferred from the intersection of mutations. If a set of mutations are co-existed in the ancestral cell and the absence of mutations in the daughter cells are more likely caused by false detection in single cell libraries or genetic draft during cell replications. Then the observed cells with different intersections (e.g *V1+V2*) will be as expected by *P_v1_*P_v2_*N*, after normalized by sequencing depth. The exclusive of intersections from high-frequency mutations will infer the separation of mtDNA mutations and multiple cell lineage. The intersections of the variants were quantified by the Upset R package^11^. In the scATAC-seq from pHSCs from SU353, the intersection of variants showed most of the cells were separated by two sets of different variants (**Fig. 2d**). But there are a few cells displaying a mixture of variants from the two sets. We suspected these may cause by the doublet of cells in the same well during single cell separation on C1 chip. We further separated the intersection map by the chip and observed the number of cells with mixture variants correlated to the concentration of cells loaded to C1 Chip (**Supplementary Figs.8**). These cells were removed during subsequent analysis. Single cells with any variants in the two sets were kept and cells with more than 40X coverage on mtDNA, but no variants in the two sets were considered as wild-type HSCs. After all the filter steps, 153 cells had lineage information and were separated into three subpopulations.

**Single cell ATAC-seq chromatin analysis.** ATAC sequences mapped to the nuclear genome were used for chromatin accessibility profiling. Bam files were merged for the same cell types and used as input files for chromVAR^14^. Peak files from Buenrostro et.al^4^ were used as open background regions to quantify the accessibility signal from every single cell. Cells with fewer than 200 unique reads or less than 25% of reads in peak regions were removed for chromatin analysis. chromVAR was applied to calculate TF motif-associated chromatin accessibility landscape changes and identify potential regulators of epigenomic variability. This approach quantifies accessibility variation across single-cells by aggregating accessible regions containing a specific TF motif, then compares the observed accessibility of all peaks containing a TF motif to a background set of peaks normalizing for known technical confounders. For determining differentially accessible motifs between different subpopulations, a Wilcoxon test was used to calculate the p values of the difference between the two groups.

**Molecular Phylogenetic analysis by Maximum Likelihood method.** The mitochondrial sequence for each individual was reconstructed according to the homoplasmic variants and the reference sequence (rCRS). Mitochondrial sequences from 11 patients were formatted as the input for MEGA(6.06)^15^ for phylogenetic analysis. Sequences were imported as nucleotide sequences. The evolutionary history was inferred by using the Maximum Likelihood method based on the Tamura-Nei model*.* The tree with the highest log likelihood (-23855.2085) is shown. Initial tree(s) for the heuristic search were obtained automatically by applying Neighbor-Join and BioNJ algorithms to a matrix of pairwise distances estimated using the Maximum Composite Likelihood (MCL) approach, and then selecting the topology with superior log likelihood value. The tree is drawn to scale, with branch lengths measured in the number of substitutions per site.

**Code availability.** Custom analysis code can be downloaded from GitHub(https://github.com/ChangLab/ATAC_mito_sc)

11.	Conway, J. R., Lex, A. & Gehlenborg, N. UpSetR: an R package for the visualization of intersecting sets and their properties. doi:10.1093/bioinformatics/btx364

12.	Coller, H. A. *et al.* High frequency of homoplasmic mitochondrial DNA mutations in human tumors can be explained without selection. *Nat. Genet.* **28,** 147–150 (2001).

13.	Miller, F. J., Rosenfeldt, F. L., Zhang, C., Linnane, A. W. & Nagley, P. Precise determination of mitochondrial DNA copy number in human skeletal and cardiac muscle by a PCR-based assay: lack of change of copy number with age. *Nucleic Acids Res.* **31,** e61 (2003).

14.	Schep, A. N., Wu, B., Buenrostro, J. D. & Greenleaf, W. J. chromVAR: inferring transcription-factor-associated accessibility from single-cell epigenomic data. *Nat. Methods* **14,** 975–978 (2017).

15.	Tamura, K., Stecher, G., Peterson, D., Filipski, A. & Kumar, S. MEGA6: Molecular Evolutionary Genetics Analysis Version 6.0. doi:10.1093/molbev/mst197
